# Electrophysiological signatures of hierarchical learning

**DOI:** 10.1101/2021.03.09.434666

**Authors:** Meng Liu, Wenshan Dong, Shaozheng Qin, Tom Verguts, Qi Chen

## Abstract

Human perception and learning is thought to rely on a hierarchical generative model that is continuously updated via precision-weighted prediction errors (pwPEs). However, the neural basis of such cognitive process and how it unfolds during decision making, remain poorly understood. To investigate this question, we combined a hierarchical Bayesian model (i.e., Hierarchical Gaussian Filter, HGF) with electrophysiological (EEG) recording, while participants performed a probabilistic reversal learning task in alternatingly stable and volatile environments. Behaviorally, the HGF fitted significantly better than two control, non-hierarchical, models. Neurally, low-level and high-level pwPEs were independently encoded by the P300 component. Low-level pwPEs were reflected in the theta (4-8 Hz) frequency band, but high-level pwPEs were not. Furthermore, the expressions of high-level pwPEs were stronger for participants with better HGF fit. These results indicate that the brain employs hierarchical learning, and encodes both low- and high-level learning signals separately and adaptively.

## Introduction

Humans and other agents need to learn about the complexity of the world in order to act adaptively. However, our inability to directly observe many important aspects of the world, makes accurate learning difficult. To learn such structure, predictive coding theories postulate that the brain uses a hierarchical generative model to infer the environmental causes of its sensory data. Crucially, in such a generative model, the brain would use hierarchical precision-weighted prediction errors (pwPEs) to continuously update this model (Rao and Ballard 1999; Friston 2009; Deutsch 2010; Iglesias et al. 2013). Over the past few decades, functional magnetic resonance imaging (fMRI) studies have investigated the neural representation of pwPEs, and have suggested that pwPEs at different hierarchical levels may be encoded by different neural loci (Iglesias et al. 2013; Lawson et al. 2017; Powers et al. 2017; Nassar, Mcguire, et al. 2019; Deserno et al. 2020; Iglesias et al. 2021). However, the blood oxygen level dependent (BOLD) signal of fMRI measures neural activity only indirectly, and cannot disentangle the time scale and frequency bands in which these signals reside, owing to its low temporal resolution. To address these issues, we use EEG to test how humans learn and update beliefs in dynamic environments.

The notion of hierarchy is central to neurobiological accounts of brain organization. In the field of learning and decision making, several hierarchical learning models have been proposed (Botvinick et al. 2009; Mathys 2011; Ribas-Fernandes et al. 2011; Diuk et al. 2013; Mathys et al. 2014; Verbeke and Verguts 2019; Verbeke et al. 2021). These models have provided indispensable frameworks for understanding the neural substrates of learning and decision making. Mathys et al. (2011, 2014) proposed a Bayesian framework (Hierarchical Gaussian Filter, HGF) to describe a hierarchical learning process, in which different hierarchical PEs are precision-weighted and computed at different levels of an information processing cascade. Subsequently, the validity of this model was firstly verified in Iglesias et al. (2013). They conducted a probabilistic reversal learning task, a classical experimental paradigm to probe hierarchical learning processes. Here, stimulus-response contingencies switched during the experiment, and subjects had to track those contingencies. Results showed that the HGF explained the behavioral data better than nonhierarchical learning models. In this vein, a series of studies using similar computational models further revealed that this hierarchical learning model is suitable for understanding both non-social (Lawson et al. 2017; Powers et al. 2017; Nassar, Mcguire, et al. 2019; Pulcu and Browning 2019; Deserno et al. 2020; Iglesias et al. 2021) and social learning (Diaconescu et al. 2017, 2019; Cole et al. 2020; Henco et al. 2020; Sevgi et al. 2020).

EEG with excellent temporal resolution has been widely used to investigate human learning (Mars et al. 2008; Kolossa et al. 2015; Marieke et al. 2016; Stefanics et al. 2018; Nassar et al. 2019; Weber et al. 2020; Hein et al. 2021). A late, stimulus-locked positivity component called the P300, with central parietal topography, has been long known to reflect surprise (Mars et al. 2008; Kolossa et al. 2015). However, recent work suggested that it also relates to learning and belief updating (Marieke et al. 2016; Nassar et al. 2019; Hein et al. 2021). For instance, Nassar et al. (2019) conducted a predictive inference task to reveal the neural representations of learning from surprise. Results showed that larger P300 signals predicted greater learning in an environment where surprise indicated a change in environmental contingencies, but less learning in an environment where surprise was indicative of a one-off outlier.

Also Hein et al. (2021) combined the hierarchical learning model with EEG recording. They observed that lower-level pwPEs about reward tendencies were represented in the ERP signals across central and parietal electrodes peaking at 496 ms in the human brain, overlapping with the late P300 component in classical ERP analysis. However, these earlier EEG studies did not manipulate volatility, thus making it hard to disentangle lower-versus higher-level pwPEs. To address this issue, the first aim of our study is to examine whether the brain processes hierarchical pwPEs, as reflected in P300 amplitudes, and whether the encoding of lower-versus higher-level hierarchical pwPEs shows a neural dissociation.

Additionally, theta oscillations have also been strongly implicated in learning. Oscillations in mid-frontal theta frequency band (around 4–8 Hz) at around 300–450 ms after feedback onset, source-localized to the anterior cingulate cortex (ACC), are commonly elicited by external feedback (Cohen et al. 2007; Luft 2014). Previous studies suggested that these theta oscillations may represent how much control should be exerted (Verbeke and Verguts 2019; Verbeke et al. 2021). Indeed, increases in frontal theta activity have been observed to relate to a wide variety of tasks under situations of cognitive control (Cohen et al. 2007; Cavanagh et al. 2010; Cavanagh and Frank 2014; Mas-Herrero and Marco-Pallarés 2014; Verbeke and Verguts 2019; Wang et al. 2020; Verbeke et al. 2021). For example, Cavanagh and Frank (2014) related theta to cognitive control by showing that theta power increased for novel and difficult stimuli, as well as for negative feedback (stimulus-locked) and errors (response-locked). Moreover, theta oscillations have been proposed to be modulated by unsigned prediction error, which is critical for understanding the neural basis of learning (Cavanagh et al. 2010). This idea has been supported by recent investigations (Mas-Herrero and Marco-Pallarés 2014; Wang et al. 2020). However, no research has directly linked theta oscillations as the feedback signals with hierarchical learning processes. Hence, this brings us to the second aim of our study: to test whether the brain processes hierarchical pwPEs reflected in theta oscillations, and whether the encoding of lower-versus higher-level hierarchical pwPEs shows a neural dissociation.

In summary, the current study aimed to apply the probabilistic reversal learning task, combining the hierarchical learning framework (HGF model) with EEG technology, to examine the cognitive and computational mechanism of hierarchical learning process in an uncertain world. We hypothesized that the brain employs hierarchical learning, and both low-and high-level pwPEs would be independently encoded by the P300 component. Additionally, we hypothesized that hierarchically related learning signals underlie theta oscillations, and the encoding of different hierarchical pwPEs may show a neural dissociation.

## Materials and Methods

### Participants

Twenty-three undergraduate students participated in the experiment for reimbursement. All of them had normal or corrected-to-normal vision, no history of hearing or neurological impairments, and were right handed. This study was approved by the Ethics Committee of the Institute of Psychology, South China Normal University. Each participant was given informed consent before the experiment. Three measured participants were not included in the analysis: one participant did not perform the task according to the instruction, and two were excluded owing to excessive EEG artifacts. Accordingly, the final sample included twenty subjects (8 females), age from 19 to 24 years old (*M* = 22.12 ± 1.3 years).

### Experimental procedure

Participants were comfortably seated in an electrically shielded room at a distance of 90 cm from the computer screen. Visual stimuli were presented at the center of the screen. Each participant completed a probabilistic reversal learning task, where participants had to learn the predictive strength of cues (animal or clothing) and predict a subsequent visual stimulus (horizontal or vertical grating) (see Figure 1A). Each type of cues consisted of one of two pictures (i.e., four pictures in total, each one was presented 25%). These images were selected from Zhang and Yang’s photo gallery (Zhang and Yang 2003; Liu et al. 2011; Wang and Zhang 2021). Critically, in our task the cue-outcome association strength changed over time. In the first stable environment, the probability of horizontal given an animal cue was 80%; in the first volatile environment, the probability of horizontal given an animal switched between 10% and 90%; in the second stable environment, the probability of horizontal given an animal was 20%; and finally, in the second volatile environment, the probability of horizontal given an animal, switched between 90% and 10% (see Figure 1B). Subjects were randomly assigned to complete the task with the stable environment first (as described above), or with the volatile environment first, in order to counteract a potential block-ordering effect. In addition, the initial pairing was counterbalanced across participants. Specifically, half of the subjects underwent experimental trials where the initial pairing tendency was animal-horizontal and clothing-vertical, and the other half of the subjects underwent experimental trials where the initial pairing was animal-vertical and clothing-horizontal. Overall, there were four versions of the experimental procedure across participants; five participants used the same version.

**Figure 1.**
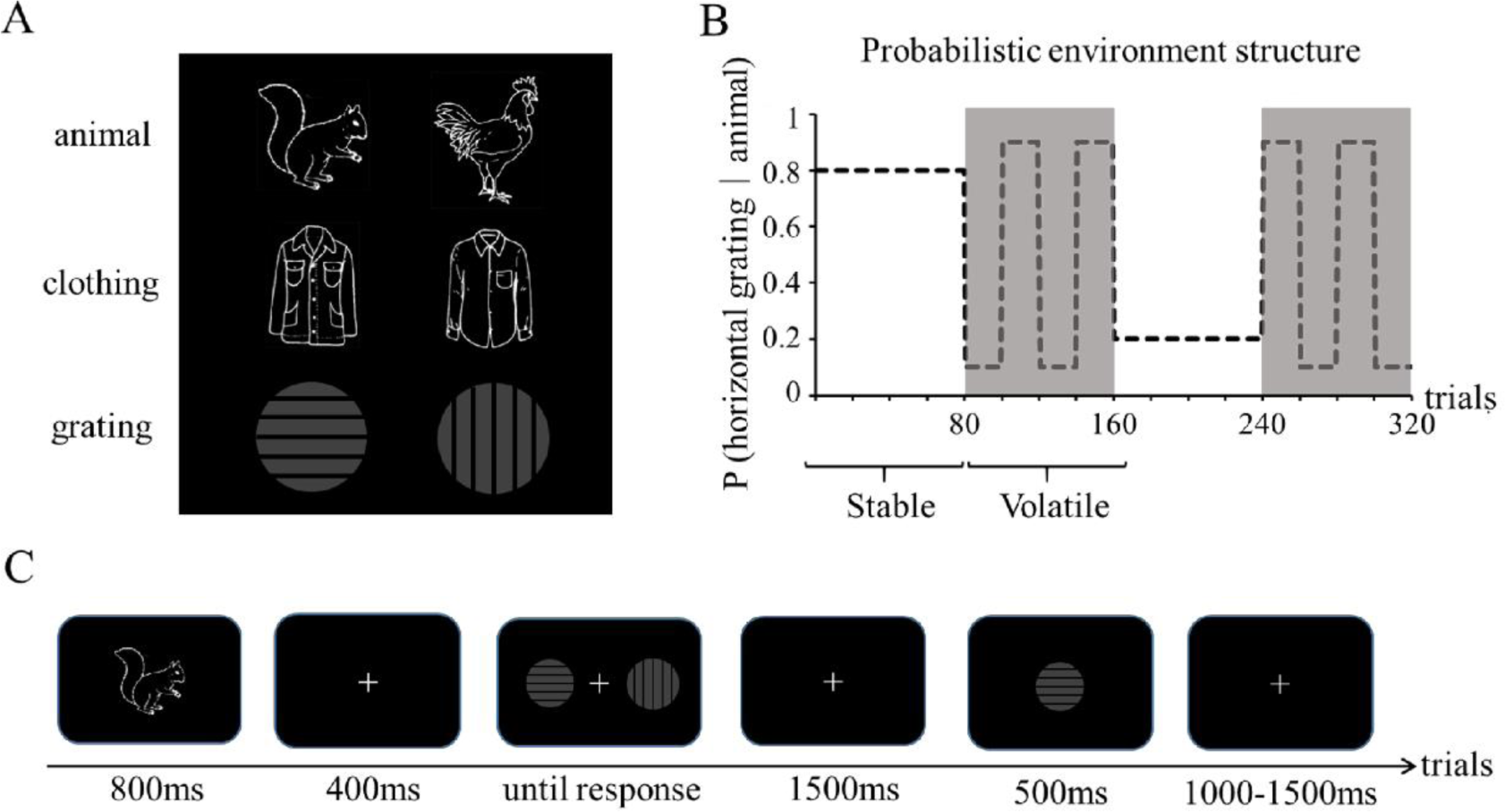
Task design. (A) Two kinds of cues (animal and clothing) and two gratings (horizontal and vertical) used in the experiment. (B) Time-varying cue-outcome contingency, in stable (white area) and volatile (gray area) environments. (C) Schematic illustration of a trial.

At the beginning of each trial, participants saw one of two possible cues (animal or clothing) for 800 ms. After a 400ms fixation cross, participants were instructed to respond as accurately and as quickly as possible to identify which of the two possible outcomes (horizontal or vertical grating) would follow, by pressing either “F” or “J”, for the left or right grating, respectively. On every trial, horizontal and vertical gratings were randomly displayed on the left and right of the screen in order to control for handedness. Subsequently, a 1000-1500ms fixation cross appeared, which was followed immediately by a 500ms feedback (i.e., a horizontal or vertical grating) (see Figure 1C). Each task began with a practice session, ensuring that participants understood the task. The entire duration of the experiment was about 40min, and participants rested for 2-3 min every 10 minutes.

### Hierarchical Gaussian Filter

To capture hierarchical learning and belief trajectories during probabilistic reversal learning, we used the Hierarchical Gaussian Filter (HGF) from Mathys et al. (2011, 2014). Software to implement the HGF was obtained from (http://www.translationalneuromodeling.org/tapas). As illustrated in Figure 2, in our task, the first level of this model represents a sequence of environmental states *x*_1_ (here: whether a horizontal or vertical grating was presented), the second level represents the cue-outcome contingency *x*_2_ (i.e., the conditional probability, in logit space, of the horizontal or vertical grating given the animal or clothing cue), and the third level the log-volatility of the environment *x*_3_. Each of these hidden states (*x*_2_ and *x*_3_) is assumed to evolve as a Gaussian random walk, such that its mean is centered around its previous value at trial *k*-1, and its variance depends on the state at the level above (see Figure 2A). Free subject-specific learning parameters were *ω_2_* (a constant component of the learning rate at the second level), *ω_3_* (the speed of learning at the third level), and *ζ* (the decision noise from the response model). Consistent with earlier studies (De Berker et al. 2016; Hein et al. 2021), we fixed the *κ* (which determined how much the estimated environmental volatility affected the learning rate at the second level) to 1. For the exact update equations at each level of the HGF model, see Mathys et al. (2011, 2014). Figure 2B illustrates the three levels of the HGF for binary outcomes and the associated belief trajectories across all 320 trials in a representative participant.

**Figure 2.**
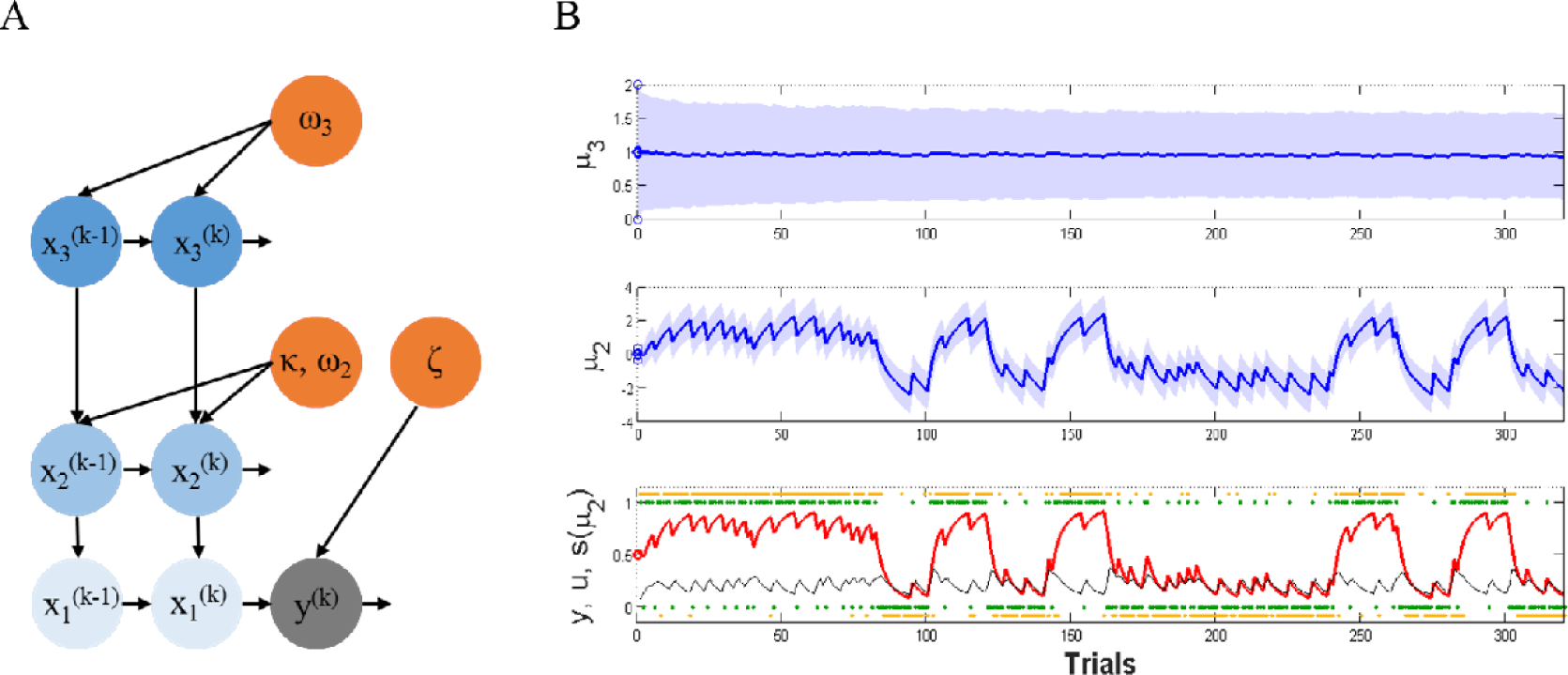
Three-level Hierarchical Gaussian Filter for binary outcomes. (A) A graphical model of the HGF with three levels. (B) Binary outcomes and model-based hierarchical belief trajectories from a representative participant. At the lowest level, *u* (green dots) correspond to the associative outcome of each trial and *y* (yellow dots) corresponded to the participant’s responses (1 = animal-horizontal or clothing-vertical, 0 = animal-vertical or clothing-horizontal). The red line of *s(μ*_2_) represented the participant’s estimate of the cue-outcome tendency *x*_2_ based on sigmoid function. The learning rate about associative outcomes at the lowest level was given in black line. At the second level, *μ*_2_ (blue line) represented the participant’s estimate of the cue-outcome tendency *x*_2_. At the top level, *μ*_3_ (blue line) represented estimates of volatility *x*_3_.

On each trial, the update equations of HGF at different levels share a general form: At any level *i* of the hierarchy, the update of the belief on trial *k* (i.e., posterior mean 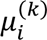of the state *x_i_*) is proportional to the pwPE 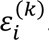. This weighted PE is the product of the PE 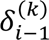 about the state on the level below and a ratio of precisions 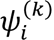

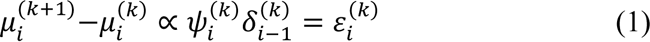

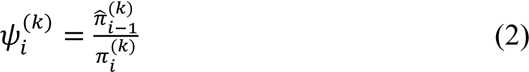

where 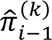 is the precision of the prediction about the level below and 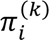 is the precision of the current belief at the current level. In fact, this precision-weighting (*ψ)* can be understood as a dynamic learning rate (Preuschoff and Bossaerts 2010; Mathys 2011; Mathys et al. 2014). The intuition behind this is that an agent’s belief updates should be more strongly driven by PEs when the precision of its data is high compared to the precision of belief in those data.

In our HGF model, two pwPEs *ε_i_* occur. At the second level, *ε*_2_ is the pwPE about the outcome of grating that serves to update the estimate of *x*_2_. At the third level, *ε_3_* is the pwPE about cue-outcome contingency that is proportional to the update of *x_3_*.

These two quantities are the focus of EEG analysis in the current study. For the specific equations, consult Mathys et al. (2011, 2014).

### Model Space

We tested three computational models of learning. The first one was a 3-level hierarchical learning model (with volatility on the third level: HGF3) explained in the previous paragraph. The second and third models are commonly used reinforcement learning models which were regarded as control models: a Rescorla Wagner (RW) model where PEs drive belief updating but with a non-adaptive learning rate (Rescorla and Wagner 1972); and a Sutton K1 model (SK1) that permits the learning rate to adapt with recent prediction errors but in a non-hierarchical way (Sutton and Barto 1998). Models were then compared at the group level for fit using random effects Bayesian model selection (BMS; Stephan et al. 2009; code from the freely available MACS toolbox; Soch and Allefeld 2018) BMS chooses the winning model based on the protected exceedance probability (PXP) which protects against the “null” possibility that there are no differences in the likelihood of models across subjects.

### EEG recording and preprocessing

EEG was recorded from a 64-channel Brain products setup, according to the modified 10-20 system, with the FCz electrode as on-line reference, and a sampling rate of 500Hz. The electrode impedance was kept below 5 kΩ. During continuous recording, a 100Hz low-pass filter was applied. Offline, EEG signals were re-referenced to averaged mastoids, and submitted to a 0.1-30Hz band-pass filter. Ocular artifact correction was performed through independent component analysis in EEGLAB (Delorme and Makeig 2004). Signals exceeding ±75 μV in any given epoch were automatically excluded. Other artifacts were removed by visual inspection.

Continuous data was segmented into -100 to 700 ms epochs around the presentation of the feedback outcome, with a pre-stimulus baseline of 100 ms. For further statistical analysis, approximately 96.2% of the data were retained after excluding artifacts.

### Time-frequency analyses

The time frequency analysis was performed to explore both phase-locked and non-phase locked brain responses elicited by feedback outcome. A windowed Fourier transform (WFT) with a fixed 200-ms Hanning window was used to obtain a time-frequency distribution (TFD) of the EEG time course. We estimated the complex time-frequency spectrum F (t, f) at each point starting from -1000 to 1500 ms (in a 2 ms interval) in latency, and from 1 to 30 Hz (in a 1-Hz interval) in the frequency domain. The spectrogram P (t, f) = |F (t, f)|^2^ represents a joint function of the power spectral density at each time frequency point (Hu et al. 2014). For each frequency, power values were baseline corrected by subtracting the mean power of the baseline (-300 to -100 ms before the feedback onset) (i.e., ER (t, f) = P (t, f) – B (f), where ER (t, f) is the event-related power value, B (f) is mean power at frequency f in the baseline) (Hu et al. 2014; Peng et al. 2019).

### Statistical analyses

For the time-domain analysis, the first GLM (GLM1) was used to test the first model-driven EEG hypothesis of a relation between P300 amplitude and hierarchical pwPEs. We selected a 0–500 ms feedback-locked interval, primarily based on the duration of the feedback (which was 500ms). This interval also covered the latency of HGF regressors in previous GLM studies (e.g., Diaconescu et al. 2017; Weber et al. 2020; although these studies used different tasks). The GLM1 used trial-wise estimates of pwPEs in level 3 (pwPE3), absolute pwPEs in level 2 (|pwPE2|), and accuracy (trial-wise correct/wrong outcome values) as regressors for the Z-scored EEG signal amplitude across all time points (0-500ms) of each electrode that was selective for feedback. Consistent with earlier studies (Iglesias et al. 2013, 2021; De Berker et al. 2016; Diaconescu et al. 2017; Hein et al. 2021), we did not orthogonalise the regressors. The absolute value for pwPE2 was selected because the direction of pwPE2 is arbitrary: the quantity *x_2_* is related to the tendency of one cue-outcome association (i.e., animal-horizontal grating). Accuracy was used as one of the regressors as we expected this variable to account for much of the signal variance in the EEG epochs. For group-level analyses, we performed one-sample *t*-tests at each single time point and electrode to examine positive or negative relations of EEG amplitudes with the trajectories of our computational quantities. In order to avoid false-positive results, an FDR correction (*q*=0.001) (Benjamini and Hochberg 1997; Benjamini and Yekutieli 2001) was applied to *p* values across all time points, electrodes, and regressors.

Additionally, we used ERP analysis to verify the results of the regression analysis. Specifically, we defined High |pwPE2|, Low |pwPE2|, High pwPE3, Low pwPE3, Positive feedback, and Negative feedback, according to trial-by-trial belief trajectory. Then, we averaged the respective EEG signal across trials, separately for each different label. Based on related studies about feedback learning (Mars et al. 2008; Kolossa et al. 2015; Marieke et al. 2016; Stefanics et al. 2018; Nassar et al. 2019; Hein et al. 2021), we analyzed the P300 components with a region-of-interest (ROI) analysis over the frontal-central scalp in the 300-500 ms range at electrodes: F1, Fz, F2, FC1, FCZ, FC2, C1, Cz, C2. The paired *t*-tests were used on the mean amplitudes with each pair of labels as a factor. Only adjusted *p*-values were reported. Furthermore, as a control analysis, we used a similar regression model (GLM2) where we replaced pwPE3 with |pwPE3| as one of regressors.

For the time-frequency domain analysis, to test the second model-driven EEG hypothesis of a relation between theta power and hierarchical pwPEs, the GLM3 was used. This contains the same regressors as described above, but as a dependent variable we used the Z-scored theta power (averaged 4-7 Hz) across all time points of each electrode that was selective for feedback. ERP analysis was also carried out to further verify the results of this regression analysis, which was similar to the time domain analysis. Here, theta power was analyzed with a region-of-interest (ROI) analysis over the frontal scalp in the 300-500 ms range and studied at electrodes: F1, Fz, F2, FC1, FCZ, FC2, according to related studies about feedback learning in the theta frequency band (Cavanagh et al. 2010; Mas-Herrero and Marco-Pallarés 2014; Wang et al. 2020). Paired *t*-tests were used on the mean power with each pair of labels as factors. Only adjusted *p*-values were reported. Furthermore, as a control analysis, we used a GLM4 in the theta power which was similar to GLM3, but replacing pwPE3 with |pwPE3| as one of regressors. To investigate the frequency specificity of hierarchical learning, we additionally used a similar regression model (GLM5) in the alpha frequency band (averaged 8-13 Hz), to investigate the relationship between hierarchical learning and alpha power.

Finally, to investigate individual differences in hierarchical learning, we tested whether the BIC values of the HGF model were related to the behavioral and EEG indices. Here, we used Pearson correlation coefficient to calculate the correlations between: (a) model-free behavioral indices and BIC; (b) hierarchical LRs and BIC; (c) hierarchical pwPEs and BIC; (d) hierarchical pwPEs effects (β values for hierarchical pwPEs effects in GLM analyses) and BIC.

## Results

### Accuracy and response time

To analyze task performance, paired *t*-tests were performed for both accuracy and response time. Accuracy was defined as making the optimal choice (i.e., the option with highest probability of current association) on a given trial. Results demonstrated that participants made the optimal choice in 86.78% of trials in the Stable environment (*SD* = 0.016), which was significantly more than in the Volatile environment (*M* = 82.06%, *SD* = 0.013), *t* (1, 19) = 5.028, *p* < 0.001 (see Figure 3A). The RT in the Stable environment (*M* = 834.71, *SD* = 65.13) was not significantly different from that in the Volatile environment (*M* = 842.76%, *SD* = 57.92), *t* (1, 19) = -0.295, *p* = 0.771 (see Figure 3B).

**Figure 3.**
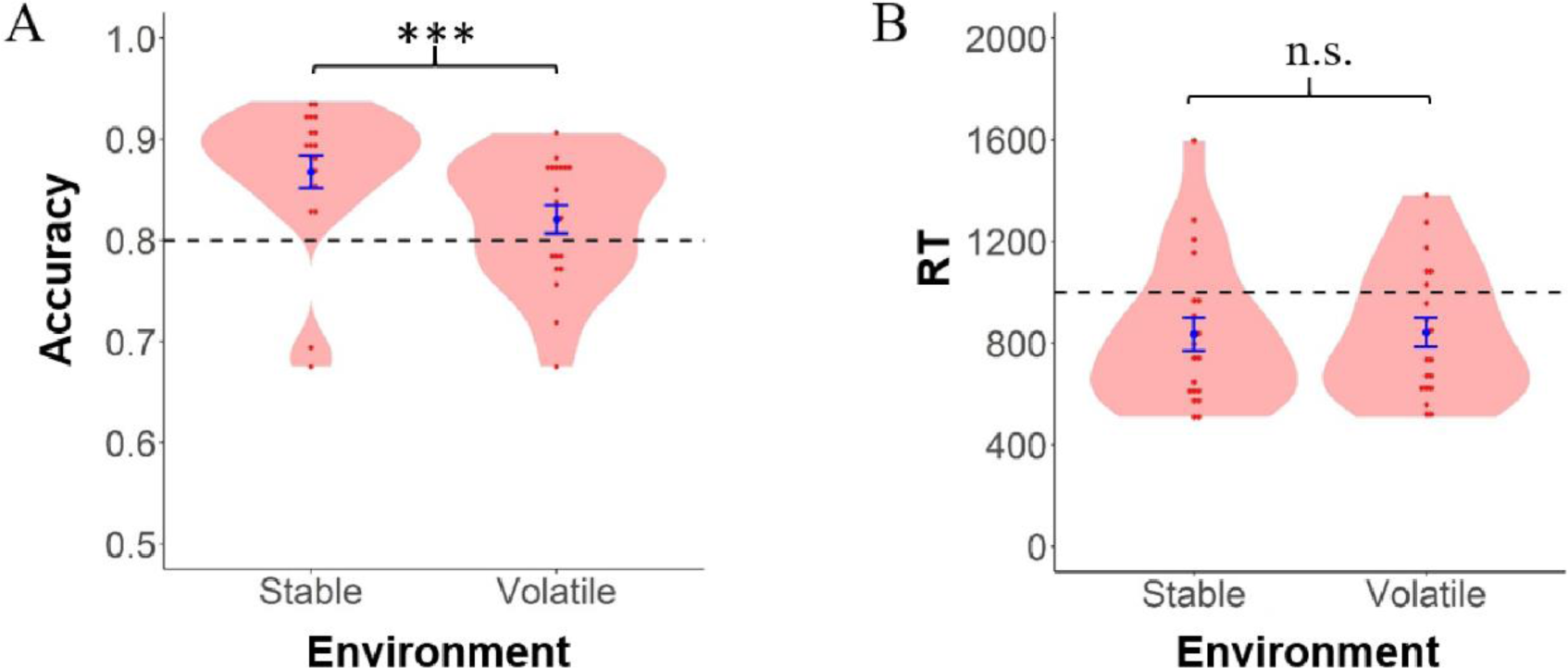
Accuracy and response time. (A) Mean accuracy for the Stable and Volatile environment. (B) Mean response time for the Stable and Volatile environment. Note: \*\*\**p* < 0.001.

### Bayesian model comparison and selection

We fitted each model individually to each of the 20 participants and obtained log-model evidence (LME), Akaike information criterion (AIC), and Bayesian information criterion (BIC) values for each. For 17 out of 20 people, the HGF3 model fitted better (smaller BIC), and for 3 out of 20 people, the RW model fitted better (smaller BIC). In addition, Bayesian model comparisons were applied to select the model with the highest protected exceedance probability (PXP). Comparisons of the three models showed that the HGF3 model was clearly superior to the other two alternative models: the results of Bayesian model selection are summarized in Table 1. Consequently, all subsequent analysis was based on the winning model only (HGF3).

**Table 1.** Bayesian model selection.

|  | HGF3 | SK1 | RW |
| --- | --- | --- | --- |
| LME | $-119.938 \pm 41.523$ | $-127.684 \pm 42.353$ | $-128.779 \pm 52.697$ |
| AIC | $223.817 \pm 80.995$ | $237.675 \pm 81.708$ | $230.658 \pm 80.916$ |
| BIC | $235.122 \pm 80.995$ | $252.748 \pm 81.708$ | $238.431 \pm 81.494$ |
| EXP | 0.909 | 0.045 | 0.046 |
| XP | 1.000 | 0.000 | 0.000 |
| PXP | 1.000 | 0.000 | 0.000 |

Note:
<sup>a</sup> HGF3, hierarchical Gaussian filter with 3 levels; SK1, Sutton K1 model; RW, Rescorla Wagner model.
<sup>b</sup> LME, log-model evidence; AIC, Akaike information criterion; BIC, Bayesian information criterion; EXPs, expected posterior probabilities; XPs, exceedance probabilities; PXPs, protected exceedance probabilities.

### Learning rate and prediction error

The probability (i.e., 2^nd^ level) learning rate (averaged across trials) was significantly higher in the Volatile environment (*M* = 1.479, *SD* = 0.281) than in the Stable environment (*M* = 1.393, *SD* = 0.239), *t* (1, 19) = -8.084, *p* < 0.001 (see Figure 4A). The volatility (i.e., 3^rd^ level) learning rate in the Stable environment (*M* = 0.040, *SD* = 0.0067) was not significantly different from that in the Volatile environment (*M* = 0.039, *SD* = 0.0062), *t* (1, 19) = 0.224, *p* = 0.825 (see Figure 4B).

**Figure 4.**
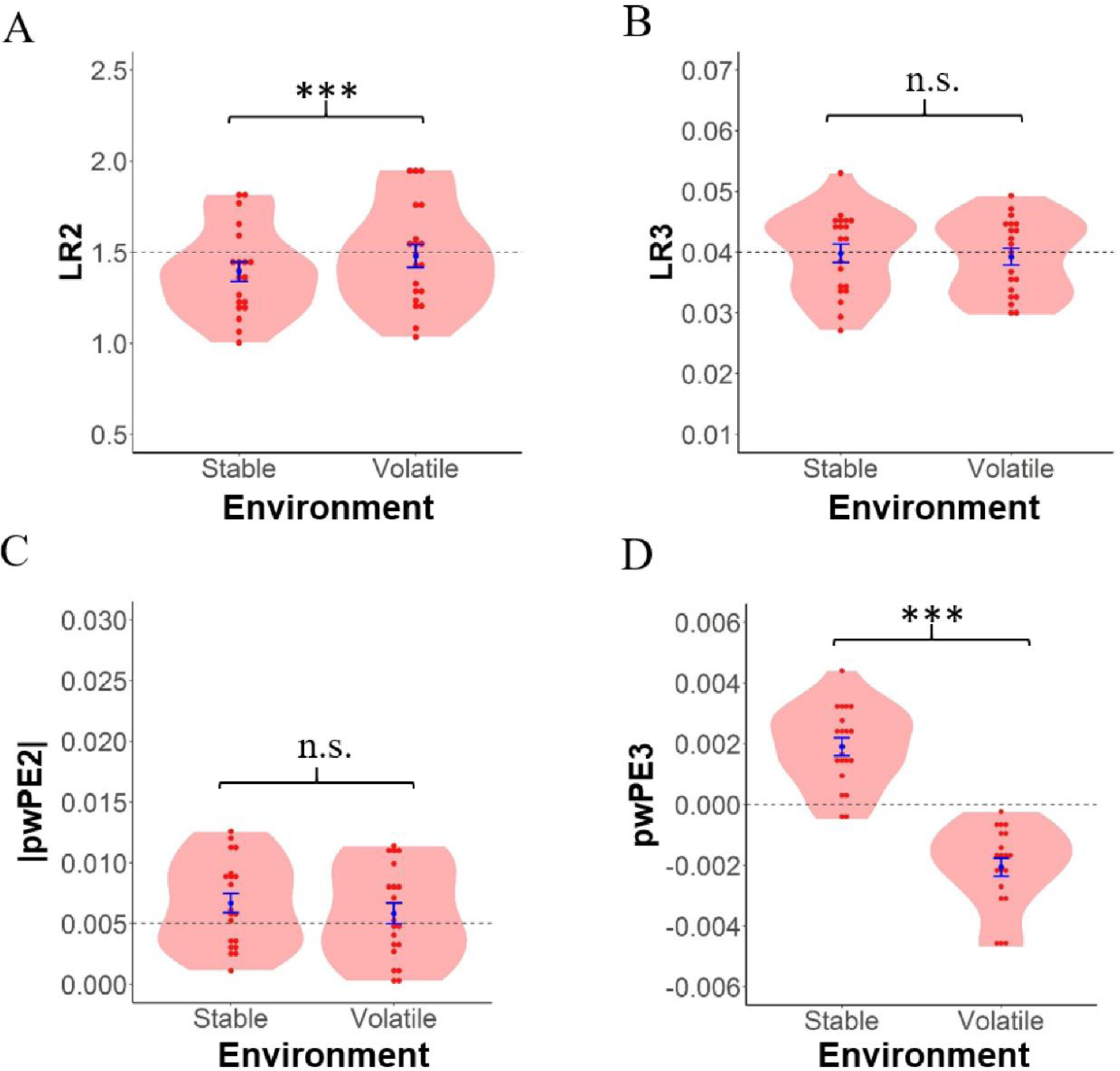
Learning rate and prediction error. Subject-specific (A) probability learning rates (*ψ_2_*), (B) volatility learning rates (*ψ_3_*), (C) probability precision-weighted prediction errors (*ε_2_*), and (D) volatility precision-weighted prediction errors (*ε_3_*) for the Stable and Volatile environment, respectively. Note: \*\*\**p* < 0.001.

The absolute probability (i.e., 2^nd^ level) pwPE (averaged across trials) in the Stable environment (*M* = 0.0064, *SD* = 0.0038) was not significantly different from Volatile environment (*M* = 0.0058, *SD* = 0.0038), *t* (1, 19) = 0.721, *p* = 0.479 (see Figure 4C). The volatility (i.e., 3^rd^ level) pwPE in the Stable environment (*M* = 0.0019, *SD* = 0.0013), was significantly higher than in the Volatile environment (*M* = -0.0021, *SD* = 0.0013), *t* (1, 19) = 7.925, *p* < 0.001 (see Figure 4D).

### Single-trial amplitude modulated by hierarchical precision-weighted PEs

**Effect of |pwPE2|:** In a dynamic learning environment, |pwPE2| significantly modulated trial-wise ERP responses from 342ms to 500ms post feedback stimulus over frontal and central electrodes (*P_FDR_* < 0.0001). Consistently, statistical analysis of the P300 component (300-500ms) over the frontal-central scalp revealed that a high |pwPE2| elicited larger positivity than low |pwPE2|, *t* (1, 19) = 15.084, *p* < 0.001 (see Figure 5A).

**Figure 5.**
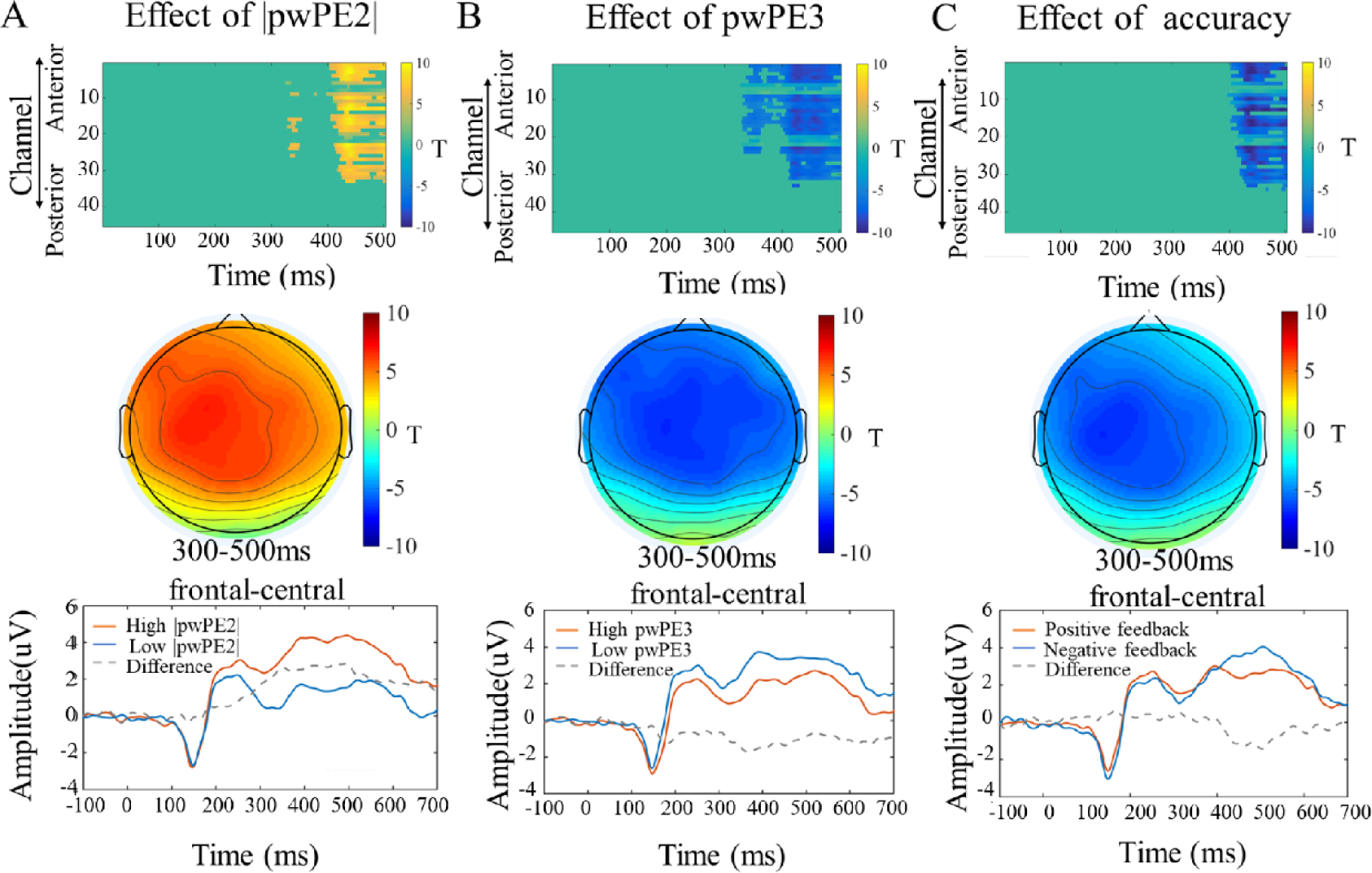
Signatures of (feedback-locked) precision-weighted prediction errors and feedback outcome on trial-wise ERPs. (A) The trajectories of |pwPE2| significantly modulated trial-wise ERPs responses from 342ms to 500ms post feedback stimulus over frontal and central electrodes. (B) The trajectories of pwPE3 significantly modulated trial-wise ERPs responses from 330ms to 500ms post feedback stimulus over frontal and central electrodes. (C) The trajectories of accuracy significantly modulated trial-wise ERPs responses from 417ms to 500ms post feedback stimulus over frontal and central electrodes. Note: *P_FDR_* < 0.0001.

**Effect of pwPE3:** In the same regression model as pwPE2 (i.e., GLM1), results revealed that pwPE3 significantly modulated trial-wise ERP responses from 330ms to 500ms post feedback stimulus over frontal and central electrodes (*P_FDR_* < 0.0001).

Consistently, statistical analysis of the P300 component (300-500ms) over frontal and central electrodes revealed that the low pwPE3 elicited larger positivity than high pwPE3, *t* (1, 19) = -8.084, *p* < 0.01 (see Figure 5B).

**Effect of accuracy:** Regression model results revealed that accuracy significantly modulated trial-wise ERP responses from 417ms to 500ms post feedback stimulus over frontal and central electrodes (*P_FDR_* < 0.0001). Consistently, statistical analysis of the P300 component (400-500ms) over frontal and central electrodes revealed that negative feedback elicited a larger positivity than positive feedback, *t* (1, 19) = 13.624, *p* < 0.001 (see Figure 5C).

As a control analysis, we additionally used a similar regression model (GLM2) in the ERP amplitudes, where we replaced pwPE3 with |pwPE3| as one of regressors.

However, no significant results was reported for any of the regressions (*P_FDR_* < 0.001).

In sum, we found a linear relationship between post-feedback ERP responses and specific hierarchical pwPEs.

### Single-trial theta frequency power modulated by hierarchical precision-weighted PEs

**Effect of |pwPE2|:** We next investigated GLM3. Here, |pwPE2| significantly modulated trial-wise theta power responses from 307ms to 500ms post feedback stimulus over frontal electrodes (*P_FDR_* < 0.001). Furthermore, statistical analysis of the theta power (300-500ms) over frontal electrodes revealed that the high |pwPE2| elicited more theta power relative to low |pwPE2|, *t* (1, 19) = 13.621, *p* < 0.001 (see Figure 6A).

**Figure 6.**
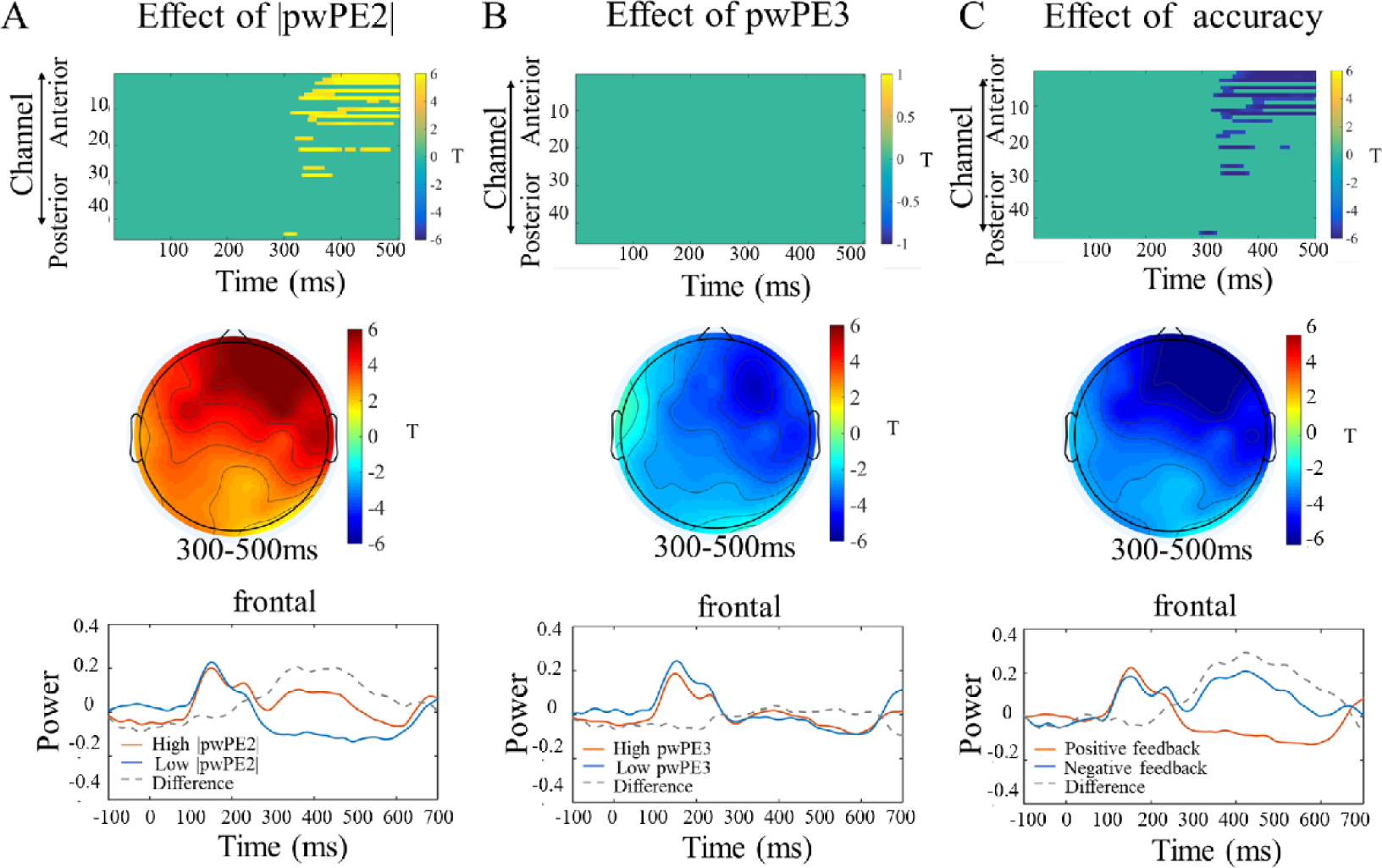
Signatures of (feedback-locked) precision-weighted prediction errors and feedback outcome on trial-wise theta frequency power. (A) The trajectories of |pwPE2| significantly modulated trial-wise theta power responses from 307ms to 500ms post feedback stimulus over frontal electrodes. (B) The trajectories of pwPE3 did not significantly modulate the theta power responses. (C) The trajectories of accuracy significantly modulated trial-wise theta power responses from 306ms to 500ms post feedback stimulus over frontal electrodes. Note: *P_FDR_* < 0.001.

**Figure 7.**
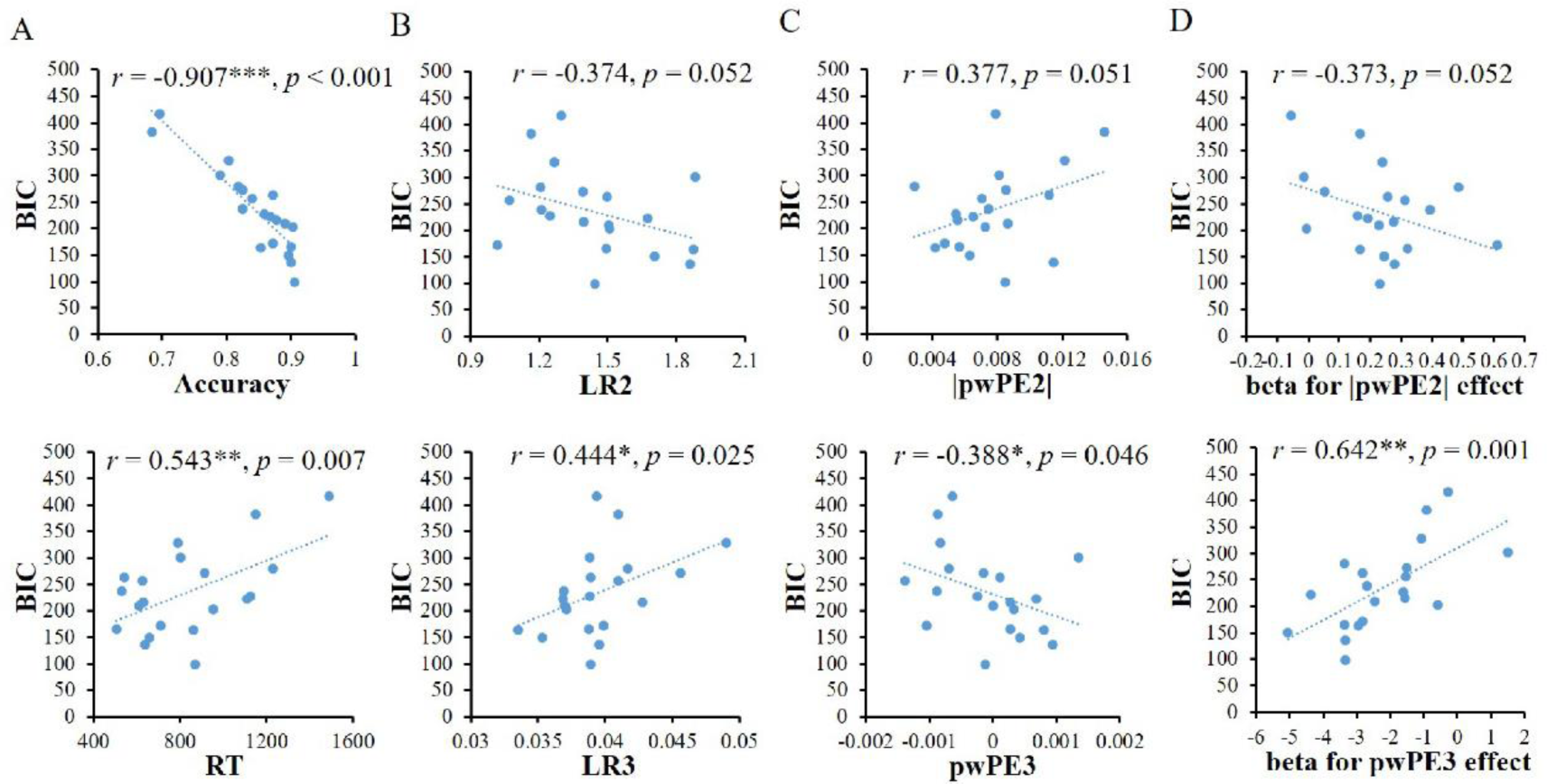
Correlations between behavioral or EEG indices and BIC of the HGF model. (A) Model-free behavioral indices and BIC. (B) Hierarchical RTs and BIC. (C) Hierarchical pwPEs and BIC. (D) Hierarchical pwPEs effects and BIC. Note: \*\*\**p* < 0.001; \*\**p* < 0.01; \**p* < 0.05.

**Effect of pwPE3:** Results from the same regression model (GLM3) revealed that pwPE3 did not significantly modulate the theta power responses. Besides, statistical analysis of the theta power (300-500ms) over frontal electrodes revealed that theta power in the high pwPE3 condition was not significantly different from low pwPE3 condition, *p* = 0.771 (see Figure 6B).

**Effect of accuracy:** Regression model results revealed that accuracy significantly modulated trial-wise theta power responses from 306ms to 500ms post feedback stimulus over frontal electrodes (*P_FDR_* < 0.001). Next, statistical analysis of the theta power (300-500ms) over frontal electrodes revealed that the negative feedback elicited more theta power than positive feedback, *t* (1, 19) = 18.762, *p* < 0.001 (see Figure 6C).

As a control analysis, we used a similar regression model (GLM4) for theta power, where we replaced pwPE3 with |pwPE3| as one of regressors. However, no significant results were found for any of the regressors (*P_FDR_* < 0.001).

In sum, we found a linear relationship between post-feedback theta power and low-level but not with high-level pwPEs.

### Single-trial alpha frequency power modulated by hierarchical precision-weighted PEs

For exploratory purposes, we used the same regression model in the alpha frequency band (averaged 8-13 Hz) (GLM5) to investigate the relationship between hierarchical learning and alpha power. However, no significant result was reported for any of the regressions (*P_FDR_* < 0.001).

***Individual differences in model fits in relation to the behavioral and EEG indices*** For the model-free behavioral indices, a correlation test across subjects between accuracy and BIC revealed that the HGF model fitted significantly better for participants with a higher accuracy (*r* = -0.907, *p* < 0.001). Additionally, there was a significant positive correlation between RT and BIC (*r* = 0.543, *p* = 0.007) (see Figure

9A). As a control analysis, we calculated the correlation between model-free behavioral indices (accuracy, RT), and BIC of the other models. Participants with better performance had a better RW model fit (for accuracy, *r* = -0.886, *p* < 0.001; for RT, *r* = 0.538, *p* = 0.007). A similar result was obtained for the SK1 model (for accuracy, *r* = -0.874, *p* < 0.001; for RT, *r* = 0.545, *p* = 0.006).

For the model-based behavioral indices, a correlation test between LR3 and BIC revealed that the HGF model fitted significantly better for participants with a lower LR3 (*r* = 0.444, *p* = 0.025). However, the analysis showed a trend of a negative correlation between LR2 and BIC (*r* = -0.374, *p* = 0.052) (see Figure 9B).

Additionally, a correlation test between |pwPE2| and BIC only showed a trend of a positive correlation (*r* = 0.377, *p* = 0.051). However, the analysis revealed that the HGF model fitted significantly better for participants with a higher pwPE3 (*r* = - 0.388, *p* = 0.046) (see Figure 9C).

For the time domain indices, we extracted the β values of each regressor and each participant and calculated the correlation between BIC values and β values for each regressor. As can be seen in the figures 9D, the |pwPE2| effect was stronger for participants with better behavioral fit (lower BIC) of the HGF model (the correlation between |pwPE2| effect and BIC values was marginally significant, *r* = -0.373, *p* = 0.052). Furthermore, the pwPE3 effect was stronger for participants with better HGF fit (the correlation between pwPE3 effect and BIC values was significant, *r* = 0.642, *p* = 0.001) (see Figure 9D). However, for the time-frequency domain indices, both the low-level and high-level pwPE effect did not differ significantly for BIC values (*p* > 0.1).

## Discussion

The current study combined HGF model and EEG technology to examine how the brain processes hierarchical learning signals at the electrophysiological level.

Behaviorally, the HGF fitted the observed behavioral data much better than two alternative non-hierarchical models. This suggests that participants were indeed likely to engage in hierarchical learning. Neurally, our results supported the hypothesis that the temporal dynamics of hierarchical learning signals were reflected in trial-by-trial changed ERP amplitude. More specifically, we found that both low- and high-level pwPE, independent of one another (i.e., in the same regression model), were encoded by the P300 component. Moreover, the trajectories of low-level but not high-level pwPEs were associated with increases in theta power, again indicating that different hierarchical learning signal may be separately represented in the brain. Furthermore, the expressions of high-level pwPEs were stronger for participants with better HGF fit, suggesting that there are individual differences in the extent to which hierarchical learning is applied in the probabilistic reversal learning task.

The empirical advantage of the HGF in the present study was in line with previous studies proposing that a hierarchical learning framework is suitable for understanding learning (Mathys 2011; Iglesias et al. 2013, 2021; Mathys et al. 2014). In this model, the low-level (i.e., 2^nd^ level) learning rates were significantly higher in the volatile environment than in the stable environment. This matches previous findings about adaptive learning (Behrens et al. 2007, 2008; Silvetti et al. 2013; Browning et al. 2015; D’Acremont and Bossaerts 2016; Nassar et al. 2019; Gagne et al. 2020), which demonstrated human learning behavior can adapt to the changes of environment. For example, Behrens et al. (2007) firstly tested whether manipulations of environmental volatility can alter humans’ learning rates. They proposed that volatile environments required increased learning in the face of surprising information, while stable environments dictated less learning from surprising information. Consistently, our findings suggest that hierarchical learning rates (i.e., 2^nd^ level) are adjusted adaptively according to changes in a dynamic environment.

According to the HGF model, the low-level learning rate about cue-outcome probabilities adaptively adjusts the weighting of prior expectations and sensory inputs (Mathys 2011; Mathys et al. 2014). Accordingly, people showed a higher low-level learning rate (i.e., more weighting of sensory input) in the volatile compared to stable environment.

Our present results provided another behavioral piece of evidence supporting an adaptive learning mode in a dynamic environment. Specifically, in our data the high-level (i.e., 3rd level) pwPEs in the stable environment were significantly higher than in the volatile environment. According to the HGF model, the high-level volatility pwPEs is proportional to the update of the belief about environmental log-volatility. If the volatility PEs is positive, this means that the environmental volatility is underestimated; conversely, it is negative if the volatility is overestimated (Mathys 2011; Mathys et al. 2014). Thus, in our data, where environments were alternatingly stable and volatile, the volatile environment was considered more changeable than it actually was relative to the stable environment, perhaps due to a contrast effect between the two conditions.

Previous EEG studies have demonstrated that the processing of learning signals is reflected in ERP amplitudes. Specifically, a central-parietal component of the P300 relates positively to learning and belief updating (Marieke et al. 2016; Nassar et al. 2019; Hein et al. 2021). Consistent with these observations, the present EEG results revealed that the P300 response on a given trial positively predicted the low-level pwPEs, but negatively predicted the high-level pwPEs. On the one hand, low-level pwPEs serves to update the estimate of the cue-outcome contingency in logit space. The higher absolute pwPE2, the stronger the belief updating in low-level cue-outcome contingency (Mathys 2011; Iglesias et al. 2013, 2021; Mathys et al. 2014; Lawson et al. 2017; Hein et al. 2021). Thus, there is a positive linear relationship between absolute pwPE2 and P300 amplitude. On the other hand, the high-level pwPEs is proportional to the update of the belief about environmental log-volatility, as described above. Therefore, the negative linear relationship between pwPE3 and P300 amplitude may suggest, that the environmental volatility was overestimated with the decrease of pwPE3, leading to learning more from environment. It is important to emphasize that both absolute pwPE2 and pwPE3 regressors survived inclusion in the multiple regression while predicting the P300 EEG signal. Thus, hierarchical belief updating (pwPEs) across different levels likely involves different cognitive processes.

Notably, time-frequency domain results revealed that theta power response on a given trial positively predicted the low-level pwPEs, but not high-level pwPEs. Those effects were specific to the theta frequency. One potential interpretation for these results can be offered in line with fMRI studies about learning in an uncertain environment (Ide et al. 2013; Iglesias et al. 2013; Diaconescu et al. 2017; Deserno et al. 2020; Henco et al. 2020) and EEG studies about theta oscillations (Cohen et al. 2007; Oliveira et al. 2007; Cavanagh et al. 2010; Luft 2014). Existing fMRI studies have linked low-level pwPEs to anterior cingulate cortex (ACC). For example, using fMRI Iglesias et al. (2013) showed that pwPEs at different hierarchical levels were encoded by different neural loci. Specifically, low-level pwPEs about stimulus probabilities were encoded by dopamine-receptive regions like DLPFC, and ACC, while high-level pwPEs about environmental volatility were encoded by the basal forebrain. Further, Ide et al. (2013) reported that participants showed activity for unsigned PEs in ACC during a Go/Nogo task. Additionally, oscillations in midfrontal theta frequency band (around 4–8 Hz), elicited after feedback onset, are source-localized to ACC (Luft 2014). Theta oscillations have been proposed to be modulated by both positive and negative (unsigned) prediction error (Oliveira et al. 2007; Cavanagh et al. 2010; Mas-Herrero and Marco-Pallarés 2014). In the same line, our results also showed modulation of absolute pwPE2 but not of pwPE2, which further supports the valence non-specificity of this effect. However, in line with the P300 results reported before (Nassar et al. 2019), we suspect that this result may be task-dependent, although this remains to be studied further.

The current research also preliminarily illustrated how individual differences in model fit can be leveraged to understand probabilistic reversal learning. Behaviorally, participants with better performance (higher accuracy or faster reaction time) had a better model fit, a result that was consistent across all three. These results suggest that participants with better performance perform in a more model-based manner, for whichever model. Here, it is of note that all models we considered (HGF3, RL, and SK1) are “model-free” models in the sense that they do not require a model of the world (Sutton and Barto 1998). It will be of interest to investigate if the current individual differences results also hold for a broader class of model-based models.

It is important to note that individual differences in (HGF3) model fit also related to the expressions of hierarchical pwPEs. More specifically, we found that the expression of pwPE3 was stronger for participants with better HGF fit. We inferred that heightened neural representation of volatility (as expressed in pwPE3) may improve hierarchical learning. Other studies have reported that a reduced expression of high-level prediction errors, could induce an overestimation of environmental volatility (Coull et al. 2011; Schmidt et al. 2012; Vinckier et al. 2016; Weber et al. 2020). Recently, Weber et al. (2020) using EEG and pharmacological manipulation (i.e., ketamine) found that the trial-by-trial relation between ERP amplitude and the high-level pwPEs was significantly diminished in the ketamine relative to the control condition. They proposed that this decreased pwPE3 disrupted the high-level inference about the environment’s statistical structure. Future research should investigate whether enhancing the expressions of pwPE3 through non-invasive brain stimulation technology may improve hierarchical learning under the probabilistic reversal learning task.

One limitation of the current study concerns the poor spatial resolution of EEG. In the present study, although EEG could record the exact time course of pwPEs, it could not reliably localize the sources of hierarchical pwPEs. Consequently, combining fMRI and EEG to investigate the hierarchical pwPEs encoding of stimulus outcome tendencies and its volatility will be an important next step. Secondly, the present study did not investigate how the brain implements communication between different brain regions under such hierarchical learning process. Oscillatory synchronization is likely a key mechanism by which neural populations transmit information and form larger networks. Future studies, with phase-based connectivity analyses (Schoffelen and Gross 2009; Rubinov and Sporns 2010; Cavanagh and Frank 2014; Verguts 2017; Wang et al. 2018; Verbeke and Verguts 2019; Verbeke et al. 2021), will be essential to reveal how the brain mechanistically coordinates neural communication to implement hierarchical learning. Finally, the current study demonstrated that hierarchical pwPEs are separately encoded in the human brain. A next relevant step will be to test whether subjects with psychiatric disorders (i.e., anxiety disorder, autism, and schizophrenia) have disturbances in one but not another hierarchical learning signal encoding. Similarly, previous hierarchical learning studies have identified that individuals with anxiety disorder, autism, or schizophrenia showed deficits in learning about a dynamic environment (Browning et al. 2015; Kaplan et al. 2016; Lawson et al. 2017; Powers et al. 2017; Pulcu and Browning 2019; Cole et al. 2020; Deserno et al. 2020). Future studies will focus on whether an abnormal encoding of hierarchical pwPEs may be related to anxiety and other psychological disorders.

To summarize, this study provided neurophysiological evidence that the brain relies on a hierarchical model of learning under dynamic environments. Hierarchical learning signals (i.e., pwPEs) at different level are separately encoded in the human brain. As such, we believe that our work furthers our understanding of hierarchical learning.

